# Reduced competition between tool action neighbors in left hemisphere stroke

**DOI:** 10.1101/547950

**Authors:** Frank E. Garcea, Harrison Stoll, Laurel J. Buxbaum

**Author notes:** **Corresponding Author**: Frank E. Garcea, Moss Rehabilitation Research Institute, Elkins Park, PA 19027.

## Abstract

When pantomiming the use of tools, patients with limb apraxia after left hemisphere stroke (LCVA) produce more spatiotemporal hand action errors with tools associated with conflicting actions for use versus grasp-to-pick-up (e.g., corkscrew) than tools having a single action for both use and grasp (e.g., hammer). There are two possible accounts for this pattern of results. Reduced performance with ‘conflict’ tools may simply reflect weakened automaticity of use action activation, which is evident only when the use and grasp actions are not redundant. Alternatively, poor use performance may reflect reduced ability of appropriate tool use actions to compete with task-inappropriate action representations. To address this issue, we developed a Stroop-like experiment in which 21 LCVA and 8 neurotypical participants performed pantomime actions in blocks containing two tools that were similar (“neighbors”) in terms of hand action or function, or unrelated on either dimension. In a congruent condition, they pantomimed the use action associated with the visually presented tool, whereas in an incongruent condition, they pantomimed the use action for the *other* tool in the block. Relative to controls and other task conditions, LCVA participants showed *reductions* in hand action errors in incongruent relative to congruent action trials; furthermore, the degree of reduction in this incongruence effect was related to the participants’ susceptibility to grasp-on-use conflict in a separate test of pantomime to the sight of tools. Support vector regression lesion-symptom mapping analyses identified the left inferior frontal gyrus, supramarginal gyrus, and superior longitudinal fasciculus as core neuroanatomical sites associated with abnormal performance on both tasks. Collectively, the results indicate that weakened activation of tool use actions in limb apraxia gives rise to reduced ability of these actions to compete for task-appropriate selection when competition arises within single tools (grasp-on-use conflict) as well as between two tools (reduced neighborhood effects).

## 1. Introduction

On a moment-to-moment basis our cognitive and neural systems need to integrate perceptual input with motor production processes in the service of object-directed action. In the study of language, vision, and memory, biased competition—that is, the weighting of inputs to enable selection among competing alternatives in accordance with task goals—is hypothesized to be a critical mechanism by which this process is achieved (e.g., see Humphreys, 1998; Kimberg & Farah, 1993; Levelt, Roelofs, & Meyer, 1999). Skilled tool use is a defining achievement of human cognition, and poses a special challenge for selection: Many tools can be used with more than one action, and many everyday tasks require more than one tool. In that context, a range of empirical and computational studies provide evidence that competition between potential actions is biased by goals and context to facilitate task-appropriate selection (e.g., see Cisek & Kalaska, 2005; Kim & Shadlen, 1999; McKinstry, Dale, & Spivey, 2008; Pastor-Bernier & Cisek, 2011; Rounis & Humphreys, 2015; for review, see Cisek & Kalaska, 2010). Perhaps surprisingly, there is relatively little research addressing the factors that influence whether particular tool actions will compete successfully with one another or the likelihood that a given tool action will be selected.

In the domains of language and semantic memory, considerable evidence suggests that the sharing of semantic (e.g., visual or propositional) or phonological features influences competition. For example, the word “robin” may interfere with the production of the word “ostrich” by virtue of the sharing of visual and propositional feature “has wings” (e.g., see Collins & Loftus, 1975; Fieder, Wartenburger, & Abdel Rahman, 2018; Mirman & Magnuson, 2008; Rose, Aristei, Melinger, & Abdel Rahman, 2018; Vieth, McMahon, & de Zubicaray, 2014; Vigliocco, Vinson, Lewis, & Garrett, 2004); as representations with shared features are more likely to compete for selection (Damian, Vigliocco, & Levelt, 2001; Luce & Pisoni, 1998; Magnuson, Dixon, Tanenhaus, & Aslin, 2007; Vigliocco et al., 2004; for review, see Chen & Mirman, 2012). Furthermore, research in patients with language deficits demonstrates that competition among neighboring word representations is exacerbated in the presence of lesions to the left inferior frontal gyrus (IFG), resulting in a greater number of selection-related errors in picture naming tasks (e.g., see Ries, Karzmark, Navarrete, Knight, & Dronkers, 2015; Schnur et al., 2009; Thompson-Schill et al., 1998). These findings are consistent with the suggestion that the left IFG provides a biasing signal that resolves competition in the selection of task-relevant lexical items (Novick, Kan, Trueswell, & Thompson-Schill, 2009; Thompson-Schill, D’Esposito, & Kan, 1999).

In the skilled action domain, eye-tracking evidence has shown that distractor tools (e.g., corkscrew) that are used with actions similar to targets (e.g., key; both used with a twisting action) cause more competition for selection than tools that are dissimilar in action (e.g., scissors), even controlling for other semantic and visual factors, and even when actions are irrelevant to the target selection task (e.g., see Lee, Middleton, Mirman, Kalenine, & Buxbaum, 2013; see also Myung, Blumstein, & Sedivy, 2006). One account of these data is that thinking about or viewing tools causes activation of actions associated with their use (e.g., see Bub & Masson, 2012), and similarity between those actions, in turn, causes the associated tools to compete with one another for selection (Buxbaum, 2017).

But what determines the similarity of action features, and thus the degree to which actions will compete? Research with non-human primates suggests that actions may be organized into ‘motor zones’ (e.g., Cooke, Taylor, Moore, & Graziano, 2003; Graziano, Cooke, Taylor, & Moore, 2004), determined in part by the ethological utility of the action (e.g., reaching, grasping, manipulating objects; for review, see Graziano, 2016). The relevance of this putative organization for competition and selection in the human brain has not, to our knowledge, been assessed. Functional magnetic resonance imaging (fMRI) research in neurotypical adults provides additional resolution on this issue, as it has been suggested that actions may be organized based on visual or motor properties, and based on features less tied to the modality through which the action stimulus was presented (e.g., the abstract function or goal of the action; inferring the intention of an actor performing an action; for review, see Lingnau & Downing, 2015; Martin, 2016). However, it remains poorly understood how these additional features contribution to the organization of action similarity and whether they compete for selection.

Our laboratory (Watson & Buxbaum, 2014) previously demonstrated that tool action “neighborhoods” are likely to be determined by at least two spatiotemporal features associated with tool use, namely, the magnitude of arm movement and the posture/movement of the hand. In a follow-up study, neurotypical participants who performed a picture-word matching task achieved lower accuracy and increased response time with items that were close versus far action neighbors, as defined by magnitude of arm movement and hand action (Watson & Buxbaum, 2014). Thus, even though action features were irrelevant to the task, close action neighbors interfered with the processing of the target stimulus. Similar effects have been observed with close semantic neighbors in language and memory domains (e.g., Wei & Schnur, 2016).

### 1.1 Abnormal Tool Use Competition in Apraxia

Across a number of experiments, research in patients with limb apraxia–a common, highly disabling, and puzzling disorder of tool use action frequently observed after left hemisphere stroke (e.g., see Johnson-Frey, 2004; Rothi, Ochipa, & Heilman, 1991) – suggests that slowed and delayed errors in tool-directed action are the result of specific, clinically significant abnormalities in the processes by which tool use actions are activated. For example, many apraxics produce substantially more action errors with tools for which there is a difference between the hand actions for using and grasping-to-move (e.g., corkscrew; hereafter ‘conflict’ tools) than with tools requiring the same action for moving and using (e.g., hammer; hereafter ‘non-conflict’ tools; e.g., Jax & Buxbaum, 2013; Watson & Buxbaum, 2015). For ‘conflict’ tools, the grasp-to-move action is closely aligned with prominent “affordances”, such that grasping processes can be driven by tools’ structural attributes, whereas tool use actions must be retrieved from memory. Moreover, the pattern of impaired performance with ‘conflict’ tools is associated with lesions to the left insula, IFG, supramarginal gyrus (SMG), and superior longitudinal fasciculus (SLF; e.g., see Watson & Buxbaum, 2015; see also Goldenberg, Hermsdorfer, Glindemann, Rorden, & Karnath, 2007; Randerath, Goldenberg, Spijkers, Li, & Hermsdorfer, 2010). Notably, despite disrupted use action representations, apraxics perform relatively normally on tasks requiring access to knowledge of tool function (i.e., purpose; Buxbaum & Saffran, 2002).

One possibility is that reduced hand action performance with ‘conflict’ tools may simply reflect weakened use action activation, which is evident only when the use and grasp actions are not redundant. Consider, for example, that grasping a cup to use it entails the same hand action as grasping to move it, such that access to the latter information may enable an apparently correct response. However, this account does not accommodate the evidence that apraxic patients exhibit abnormalities in action competition. Specifically, eye-tracking tasks have shown that individuals with apraxia exhibit *reduced* competition between tools that are used with similar hand actions (e.g., see Lee, Mirman, & Buxbaum, 2014; Myung et al., 2010). Moreover, variability in the magnitude of slow and weakened competition from tool use neighbors is correlated with the degree of overt impairment in an independent gesture production task, indicating that subtle competition abnormalities reflected in eye movements bear on deficits in overt gesture production (Lee et al., 2014). Thus, one further possibility is that reduced automaticity of tool use action activation has consequences for action competition and selection in the apraxia syndrome.

### 1.2 Motivation for the Present Study

If it is indeed the case that apraxia may be associated with abnormalities in the competitive process by which task-relevant use actions are selected, then we expect to observe relatively *good* performance by apraxic individuals in a task in which competition from automatically-activated use actions are normally disruptive. The quintessential task with those characteristics is the Stroop task. In the classic Stroop task, a prepotent response (e.g., to read words) must be inhibited in favor of a less automatically activated, task-instructed response (e.g., to name the color of the ink in which the words are printed). Using similar logic, we predicted that LCVA participants with weak and/or slowed automatic activation of use actions with familiar tool stimuli would perform relatively *well* when required to inhibit the (weakened) automatic use response and select another recently-trained use response from the same action neighborhood. Moreover, we predicted that performance on the Stroop-like task would be inversely associated with performance on a task assessing susceptibility to grasp-use conflict. Specifically, better performance on a task requiring inhibition of automatically activated use representations should be associated with more impaired performance in using ‘conflict’ tools, in which use representations must compete with grasp representations. Critically, this relationship should be specific to action; that is, no association is predicted between use of conflict tools and performance on a task requiring inhibition of tool function information.

An additional aim of the investigation was to assess the neural substrates of abnormalities in action competition. We conducted support-vector regression lesion symptom mapping analyses to assess the prediction that a core set of regions, including the left SMG and IFG, would be associated both with reduced competition on the Stroop-like task and poor performance with ‘conflict’ tools (Watson & Buxbaum, 2015; Goldenberg et al., 2007; Randerath et al., 2010).

## 2. General Methods

### 2.1 Participants

We recruited 21 chronic left cerebrovascular accident (LCVA) participants from the Neuro-Cognitive Research Registry at Moss Rehabilitation Research Institute (6 female; mean age = 61.5 years, SD = 10.1, range = 37 – 80 years; mean education = 14 years, SD = 2.1, range = 11 – 18 years). Participants were between the ages of 21 and 80, were right handed, and had suffered a single left hemisphere stroke at least 6 months prior to testing. Participants were excluded if they had a history of psychosis, drug or alcohol abuse, co-morbid neurological disorder, or severe language comprehension deficits established with the Western Aphasia Battery (Kertesz, 1982). All participants had previously been tested for with a measure of susceptibility to grasp-use conflict developed in our laboratory (see below), but were not explicitly selected based on their performance. See Table 1 for demographic information for LCVA participants. We also recruited 8 neurotypical participants from the same Research Registry (4 female; mean age = 64.7 years, SD = 9.4, range = 49-80 years; mean education = 15.9 years, SD = 1.7, range = 13 – 18 years). LCVA participants and neurotypical participants were equivalent in age (*t*(27) = 0.82, *p* = 0.42), but neurotypical participants, on average, were more highly educated (*t*(27) = 2.30, *p* = 0.03). In compliance with the guidelines of the Institutional Review Board of Einstein Healthcare Network, all participants gave informed consent and were compensated for travel expenses and participation. The informed consents obtained did not include permission to make data publicly available, as such, the conditions of our ethical approval do not permit anonymized study data to be publicly archived. To obtain access to the data, individuals should contact the corresponding author. Requests for data are assessed and approved by the Institutional Review Board of Einstein Healthcare Network.

**Table 1.**
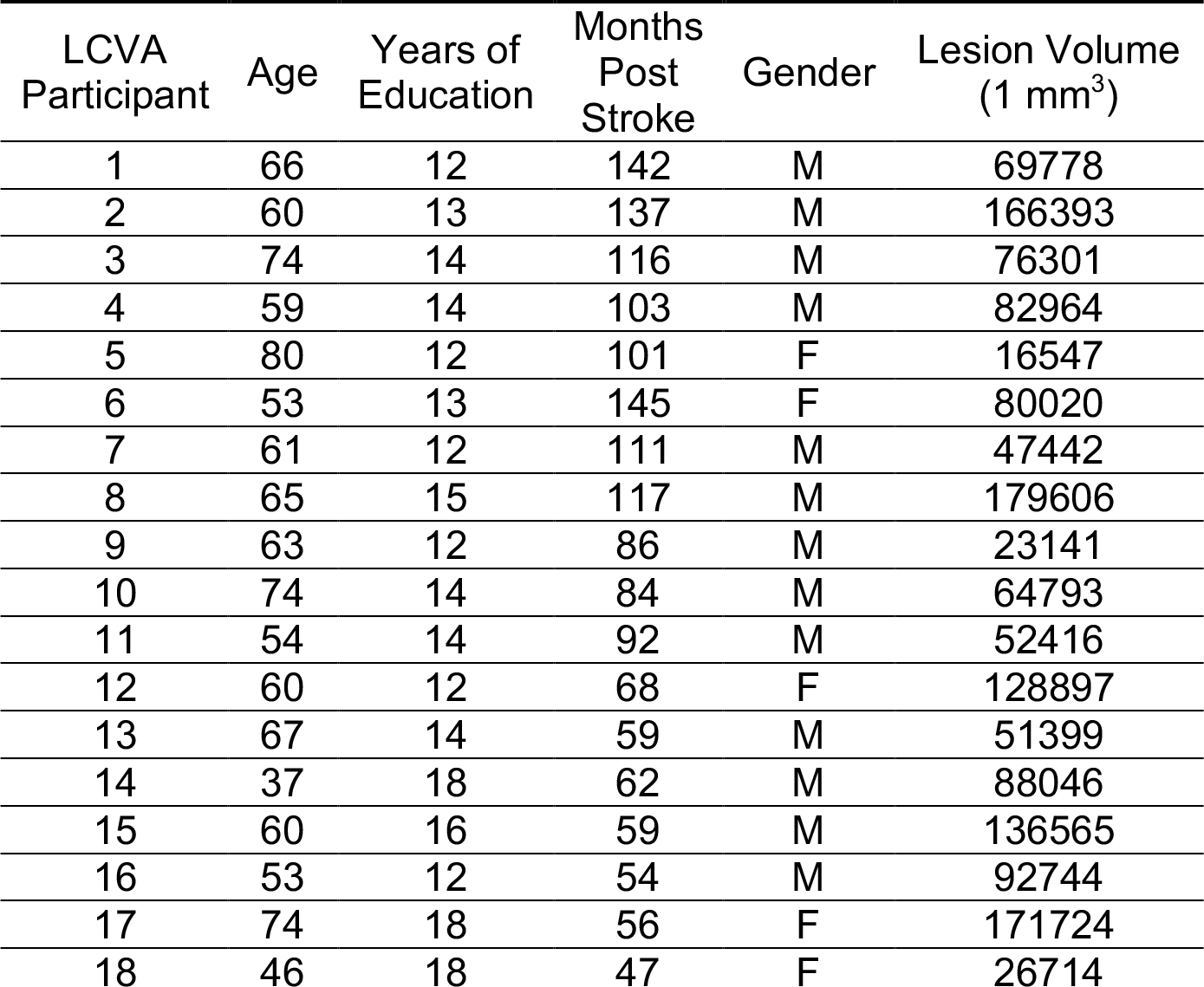
Demographic information and lesion volume for each LCVA participant.

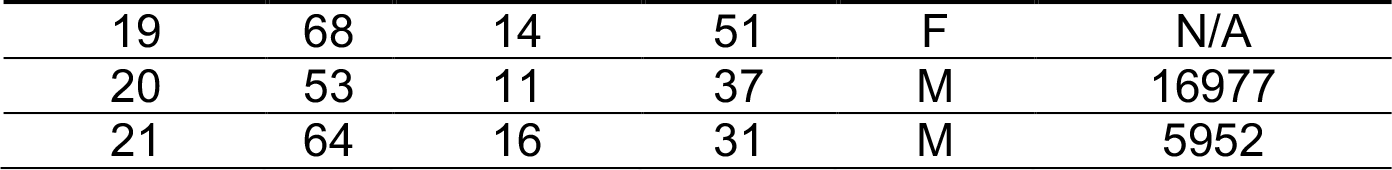

### 2.2 Neuroimaging Acquisition

Of the twenty-one LCVA individuals who participated in our study, 20 received a research-quality MRI (N = 16) or CT (N = 4) scan. MRI scans included whole-brain T1-weighted MR images collected on a 3T (Siemens Trio, Erlangen, Germany; repetition time = 1620 ms, echo time = 3.87 milliseconds, field of view = 192 × 256 mm, 1 × 1 × 1 mm voxels) or a 1.5T (Siemens Sonata, repetition time = 3,000 milliseconds, echo time = 3.54 milliseconds, field of view = 24 cm, 1.25 ×1.25 × 1.25 mm voxels) Siemens scanner, using an eight-channel head coil. Participants for whom MRI scanning was contraindicated underwent whole-brain research CT scans without contrast (60 axial slices, 3–5 mm slice thickness) on a 64-slice Siemens SOMATOM Sensation scanner.

Lesions were manually segmented on each LCVA participant’s high-resolution T1-weighted structural images. Lesioned voxels, consisting of both grey and white matter, were assigned a value of 1 and preserved voxels were assigned a value of 0. Binarized lesion masks were then registered to a standard template (Montreal Neurological Institute “Colin27”) using a symmetric diffeomorphic registration algorithm (Avants, Epstein, Grossman, & Gee, 2008, www.picsl.upenn.edu/ANTS). Volumes were first registered to an intermediate template comprised of healthy brain images acquired on the same scanner. Volumes were then mapped onto the “Colin27” template to complete the transformation into standardized space. To ensure accuracy during the transformation process, lesion maps were subsequently inspected by a neurologist (H.B. Coslett), who was naïve to the behavioral data of the study. Research CT scans were drawn directly onto the “Colin27” template by the same neurologist using MRIcron (https://www.nitrc.org/projects/mricron). For increased accuracy, the pitch of the template was rotated to approximate the slice plane of each LCVA participant’s scan. This method has been demonstrated to achieve high intra- and inter-rater reliability (e.g., see Schnur et al., 2009).

### 2.3 Experimental Task

The experimental Stroop-like task included 12 colored photographs of common tools, selected from a larger normed set of tools and manipulable artifacts from the Bank of Standardized Stimuli (BOSS) database (Brodeur, Dionne-Dostie, Montreuil, & Lepage, 2010; see also Watson & Buxbaum, 2015). Stimuli are available at https://sites.google.com/site/bosstimuli/. Based on multidimensional scaling and principal components analyses performed by Watson and Buxbaum (2014), a two-dimensional neighborhood space with “amount of arm movement” and “hand posture” ^1^ was derived, and we created 2 pairs of tools selected from that multidimensional space to be close neighbors along the hand posture dimension (corkscrew, wrench; lightbulb, key; see Watson & Buxbaum, 2014). We also created two pairs of tools that were close “function” neighbors (i.e., shared a similar purpose of use; axe, saw; matches, lighter), and two pairs that were unrelated in either action or function (fork, nail clipper; bottle opener, flyswatter).

We verified that action neighbors were similar in hand posture using ratings collected as part of a larger study measuring action similarity and tool use (see Watson & Buxbaum, 2014). Action neighbors were rated to be more similar in the hand posture required for typical use (*M*, 46; *SD*, 8.49) than function neighbors (*M*, 13; *SD*, 4.25) and unrelated items (*M*, 9; *SD*, 1.41). Watson and Buxbaum (2014) also derived a measure of semantic similarity between tools, defined as the degree to which two tools are members of the same category. Action neighbors (*M*, 1.5; *SD*, .35) and unrelated tools (*M*, 1.67; *SD*, .47) were on average less semantically similar than function neighbors (*M*, 4.75; *SD*, 0). Finally, tools were balanced for visual similarity: action neighbors (*M*, 1.38; *SD*, .18) were approximately equivalent in visual similarity to function neighbors (*M*, 1.75; *SD*, .35) and unrelated tools (*M*, 1.67; *SD*, .47). See Supplemental Table 1 for rating scores as a function of neighborhood type.

We also ensured that items were balanced for familiarity, visual complexity, and name agreement using normed values derived from the BOSS database (Brodeur et al., 2010). Critically, items were also balanced for manipulability (action neighbors, *M*, 3.3; *SD*, .37; function neighbors, *M*, 3.9; *SD*, .12; unrelated tools, *M*, 3.63; *SD*, .34), defined as the ease with which one could pantomime tool use such that a person could recognize the action. Thus, the gestures associated with items in an action neighborhood were not more difficult to pantomime or more ambiguous in the hand postures required to pantomime tool use. See Supplemental Table 2 for rating scores of each property.

**Table 2.** Average accuracy and standard error of the mean for LCVA participants and neurotypical control participants.

| Group | Action |  | Function |  | Unrelated |  |
| --- | --- | --- | --- | --- | --- | --- |
|  | Congruent | Incongruent | Congruent | Incongruent | Congruent | Incongruent |
| Neurotypical | .99 (0.04) | .94 (0.05) | .99 (0.03) | .93 (0.03) | .98 (0.03) | .92 (0.04) |
| LCVA | .80 (0.01) | .74 (0.04) | .93 (0.01) | .93 (0.03) | .92 (0.01) | .84 (0.04) |

Participants began each block with a two-portion training sequence, after which they completed the experimental task. During the first portion of training, participants were familiarized with one of the tools to appear in the experimental task (e.g., wrench and corkscrew), then shown a 400-by-400 pixel color photograph of each tool on a computer screen. They were instructed to practice pantomiming either by producing the gesture appropriate to that tool (congruent blocks) or to the other tool in the block (incongruent blocks). All responses were made with the participants’ left hands due to the possibility of right hemiparesis. The experimenter scored each gesture in real time, and upon commission of an error, provided verbal and tactile corrective feedback. Practice sequences were repeated until participants achieved a criterion of scoring 3 of 4 trials correct; upon reaching this criterion, the second portion of training began.

In the second portion of training, participants practiced a “Go/No-go” task to further familiarize them with the experimental task and stimulus sequence, and to ensure that pantomime actions were not initiated until participants were cued. Prior to the start of each trial, participants were instructed to keep their left hand relaxed and to depress a button on an E-Prime Stimulus Response Button Box; each trial started with the presentation of a 500-millisecond fixation cross, followed by presentation of a 400-by-400 pixel color photograph of a tool (a member of the same tool pair shown during the first portion of training), for 4000 milliseconds. Participants kept the button depressed until ready to gesture, whereupon they produced a pantomime and returned their finger to the button. Participants were also instructed that when an animal was presented they should not lift their finger (i.e., “no-go” trials). Each practice sequence consisted of 5 trials which included an animal image and two tools presented twice. Sequences were repeated until participants performed without error.

Following the two training sequences, the experimental sequence was conducted in two sessions, each containing 6 blocks of 28 trials (24 tool trials and 4 no-go trials) presented in random order. Figure 1 provides a schematic of the trial structure and stimuli. In each block, 2 different tools were presented, and subjects gestured either to a depicted tool (congruent blocks) or to the other tool in the block (incongruent blocks). Blocks of trials relevant to a specific type of neighborhood relation (action (A), function (F), unrelated (U)) were presented in AAFFUU-UUFFAA order for half of participants, which was reversed for the other half of participants. All participants took part in block 2 approximately one week after completing block 1. The first block of each neighborhood type was always congruent, and the second block of each neighborhood type was always incongruent. The particular tool pair shown in block one or block two was randomized across participants. Each experimental trial began with the participant depressing a button; a fixation cross was then presented for 500 milliseconds, followed by the presentation of a tool image for 4000 milliseconds. Participants responded by producing a gesture and then returning their finger to the button within the 4000-millisecond window. Across both sessions, there were 288 experimental trials (144 congruent and 144 incongruent, of which 48 each were action, function, or unrelated), and 48 no-go trials.

**Figure 1.**
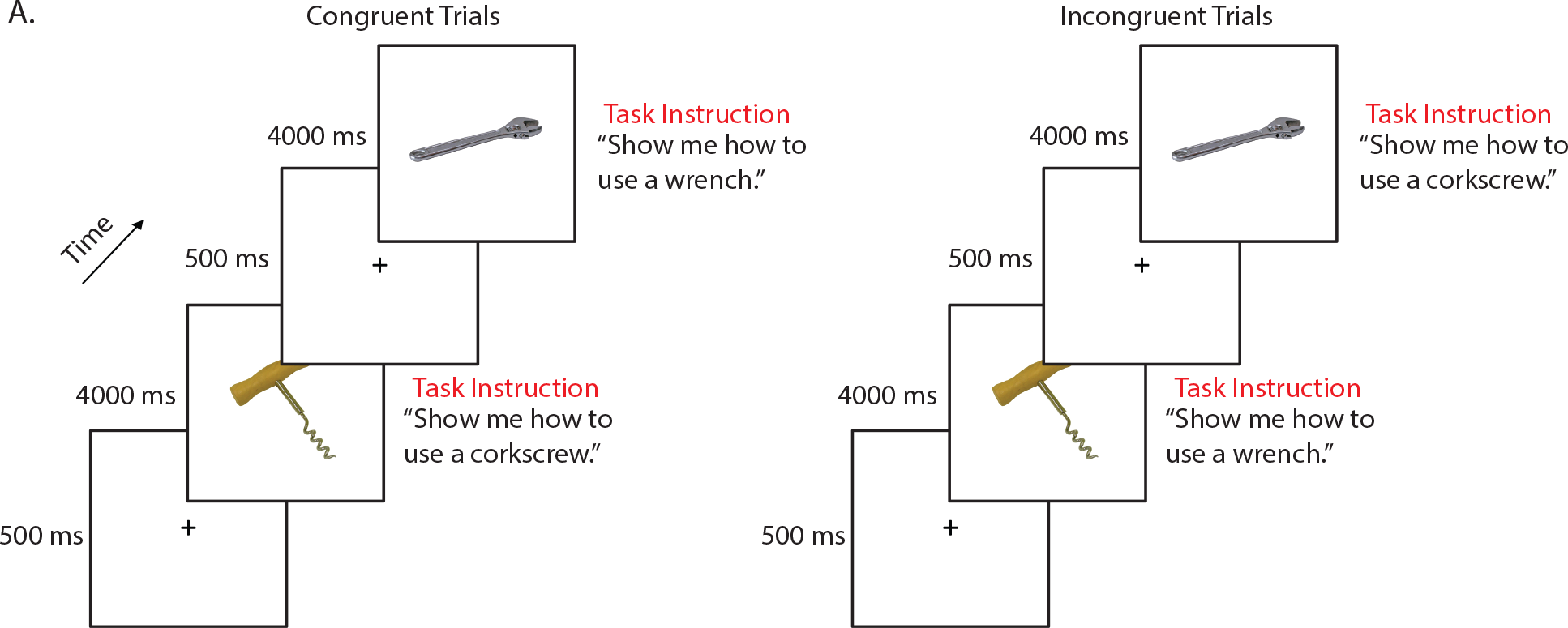
Schematic of trial structure for congruent and incongruent trials in the experimental task. Each trial began with a fixation cross presented for 500 milliseconds, followed by an image of a tool presented for 4000 milliseconds. On congruent trials participants responded by pantomiming the use of the visually presented tool; on incongruent trials participants responded by pantomiming the use of the other tool in the block.

### 2.2 Test of Grasp-Use Conflict

The test of grasp-use conflict included 40 photographs of manipulable objects (tools) taken from the BOSS database as described above. Tools included items with distinct use actions, including construction tools (e.g., wrench), household articles (e.g., teapot), office supplies (e.g., scissors), and bathroom items (e.g., razor). Half of the items had conflicting actions for use versus grasp (i.e., conflict items; e.g., corkscrew), while half of the items had a single action for both use and grasp (i.e., non-conflict items; e.g., hammer). Tools differed significantly in conflict rating (*t*(38) = 10.00, *p* < 0.001), but were equivalent in affordance strength (*t*(38)=1.50, *p* = 0.14), name agreement (*t*(38) = 1.13, *p* = 0.26), and familiarity (*t*(38) = 1.50, *p* = 0.13; for details of ratings, see Watson & Buxbaum, 2015).

Ten of the 40 tools used in the grasp-use conflict task were also included in the experimental Stroop-like task. LCVA volunteers first performed the test of grasp-use conflict, and were recruited to take part in the experimental task several months later (mean time difference, 20.6 months; SD, 13.7 months), thus mitigating the possibility of practice effects. Although we gave performance feedback during practice trials, no performance feedback was provided for either task during the experimental and grasp-use conflict tasks.

Each trial of the grasp-use conflict task began with the presentation of a 400-by-400 pixel color photograph of a tool on a computer monitor. Participants were asked to “show how you would use the tool as if you were holding and using it” with the left hand. Four practice trials with feedback (using items different than in the task itself) were given at the start of the task. As per Rothi et al. (1991), if a participant gestured the action as if their hand was the tool (body-part-as-object error), they were reminded to “show how you would use the tool as if you were actually holding it in your hand”. The first of these errors was corrected and the participant was permitted a second try.

### 2.3 Coding of Action Data

The experimental task and grasp-use conflict task were recorded with a digital camera and scored offline by two trained, reliable coders (Cohen’s Kappa score = 94%) who also demonstrated inter-rater reliability with previous coders in the Buxbaum lab (Cohen’s Kappa > 85%; see e.g., Buxbaum, Kyle, & Menon, 2005). Both tasks were coded using a portion of the detailed praxis scoring guidelines used in our previous work (see Buxbaum, Giovannetti, & Libon, 2000; Buxbaum et al., 2005; Watson & Buxbaum, 2015). First, each gesture was given credit for semantic content unless a participant performed a recognizable gesture appropriate for a semantically-related tool, including, in the case of the experimental task, the tool use gesture associated with the paired neighbor. Only gestures that were given credit for semantic content were scored on the spatiotemporal hand action dimension. A hand action error was assigned if the shape or movement trajectory of the hand and/or wrist was flagrantly incorrect, or if the hand or finger was used as part of the tool (i.e., body-part-as-object error, Buxbaum et al., 2005; Watson & Buxbaum, 2015).

### 2.4 Data Analysis Approach

Hand action scores from the experimental task were analyzed at the trial level through mixed-effect logistic regression using the R statistical programming environment version 3.5.0 with the lme4 package (Bates, Maechler, Bolker, & Walker, 2015). The R analysis code can be viewed at https://github.com/frankgarcea/GarceaStollBuxbaum. There were three main effects in our analysis: Group (LCVA participants, neurotypical participants), Neighbor Type (action, function, unrelated) and Congruence (congruent, incongruent). Prior to running each model, we assessed if there was an effect of the order of block presentation; we also included random intercepts for subject and item. All mixed-effect models were assessed by mapping the log likelihood ratio of a full and reduced model using a chi square distribution. We removed interaction terms and assessed the full model when testing for significant main effects. An alpha threshold of 0.05 was used to determine statistical significance, and all effects are reported as log odds.

Performance on the conflict versus non-conflict trials of the test of grasp-use conflict were assessed with *t*-tests. These scores were also used to derive a grasp-use conflict score, which represents hand action (HA) performance when pantomiming the use of conflict tools relative to non-conflict tools, controlling for overall performance accuracy (HA_Conflict_ − HA_Non-conflict_ / (HA_Conflict_ + HA_Non-conflict_). Negative-going grasp-use conflict scores indicate worse performance with conflict tools relative to non-conflict tools (i.e., high susceptibility to grasp-use conflict).

A final mixed-effects model assessed how competition in the test of grasp-use conflict related to performance in the experimental task. Grasp-use conflict scores and performance for incongruent action, function, and unrelated neighbor trials in the experimental task were entered as the principal dependent variables, and performance on congruent trials of the relevant condition of the experimental task served as co-variates.

### 2.5 Support Vector Regression-Lesion Symptom Mapping (SVR-LSM) Analyses

SVR-LSM was performed in MATLAB using a toolbox developed by DeMarco and Turkeltaub (2018) (https://github.com/atdemarco/svrlsmgui/). SVR-LSM is a multivariate technique that uses machine learning to determine the association between lesioned voxels and behavior when considering the lesion status of all voxels submitted to the analysis. It overcomes several limitations of voxel-based lesion symptom mapping (VLSM), including inflated false positives from correlated neighboring voxels (Pustina, Avants, Faseyitan, Medaglia, & Coslett, 2018), Type II error due to correction for multiple comparisons (Bennett, Wolford, & Miller, 2009), and uneven statistical power due to biased lesion frequency as a function of vascular anatomy (Mah, Husain, Rees, & Nachev, 2014; Sperber & Karnath, 2017). SVR-LSM has been shown to be superior to VLSM when multiple brain areas are involved in a single behavior (Herbet, Lafargue, & Duffau, 2015; Mah et al., 2014; Mirman, Zhang, Wang, Coslett, & Schwartz, 2015; for discussion, see Zhang, Kimberg, Coslett, Schwartz, & Wang, 2014).

We performed two SVR-LSM analyses. The first analysis tested the prediction that a reduced incongruence effect in the experimental task would be associated with lesions to the left insula, SMG, IFG, and SLF. The dependent measure was performance on incongruent action neighbor trials, residualized against performance on congruent action neighbor trials. Positive-going residual values indicate better performance on incongruent action neighbor trials. A second SVR-LSM analysis was performed with the Grasp-Use Conflict score, in which hand posture accuracy for conflict scores were residualized against non-conflict hand posture conflict scores. Positive-going residual values indicate a higher susceptibility to grasp-use conflict when gesturing the use of conflict tools relative to non-conflict tools, which should also be associated with lesions to the left insula, SMG, IFG, and SLF (see Watson & Buxbaum, 2015).

Only voxels lesioned in at least 15% of participants (3 participants) were included. Voxelwise statistical significance was determined using a Monte Carlo style permutation analysis (10,000 iterations) in which the critical behavioral data were randomly assigned to a lesion map, and the resulting map was set to a threshold of *p* < .05 to determine chance-level likelihood of a lesion-symptom relationship. We then further restricted the resulting map by eliminating any clusters with fewer than 500 contiguous voxels (see Lacey, Skipper-Kallal, Xing, Fama, & Turkeltaub, 2017; Skipper-Kallal, Lacey, Xing, & Turkeltaub, 2017).

The Johns-Hopkins DTI-based probabilistic white matter tractography atlas (Mori et al., 2008) was used to assess the overlap of significant voxels in the SVR-LSM analyses with major white matter fibers (e.g., superior longitudinal fasciculus) at a probability threshold of 25% (see Watson & Buxbaum, 2015).

## 3. Results

### 3.1 Excluded and Missing Trials

Experimental task trials were excluded from analysis if participants deviated from task instructions (e.g., misuse of the button box), or produced a semantic content error. For neurotypical participants, none of the trials were excluded for deviation from task instructions, and 0.6% of trials (2 trials) were excluded for semantic content errors. Two percent (125 trials) of LCVA participants’ trials were discarded due to deviation from task instructions, and 1% of trials (68 trials) were discarded due to semantic content errors; of the remaining trials, 80% (4728 trials) were correct, 15% of trials (828 trials) were hand action errors, and 5% (299 trials) were other error types (arm, amplitude) not under consideration in this study.

### 3.2 Experimental Task

Figure 2 and Table 2 present results of the experimental task. We analyzed the LCVA and neurotypical data in two separate models. The first (Model 1) contained the main effect of Group and Congruence along with a Group-by-Congruence interaction term to assess whether the groups differed in congruence effects, collapsing across neighbor type. The second (Model 2) contained the main effects of Group and Neighbor Type along with a Group-by-Neighbor Type interaction term to assess whether the groups differed in performance for different neighbor types, collapsing across congruence (a model containing an interaction term for Group-by-Neighbor type-by-Congruence failed to converge). A third model (Model 3) contained only the LCVA data, and assessed the relation between Congruence and Neighbor Type.

**Figure 2.**
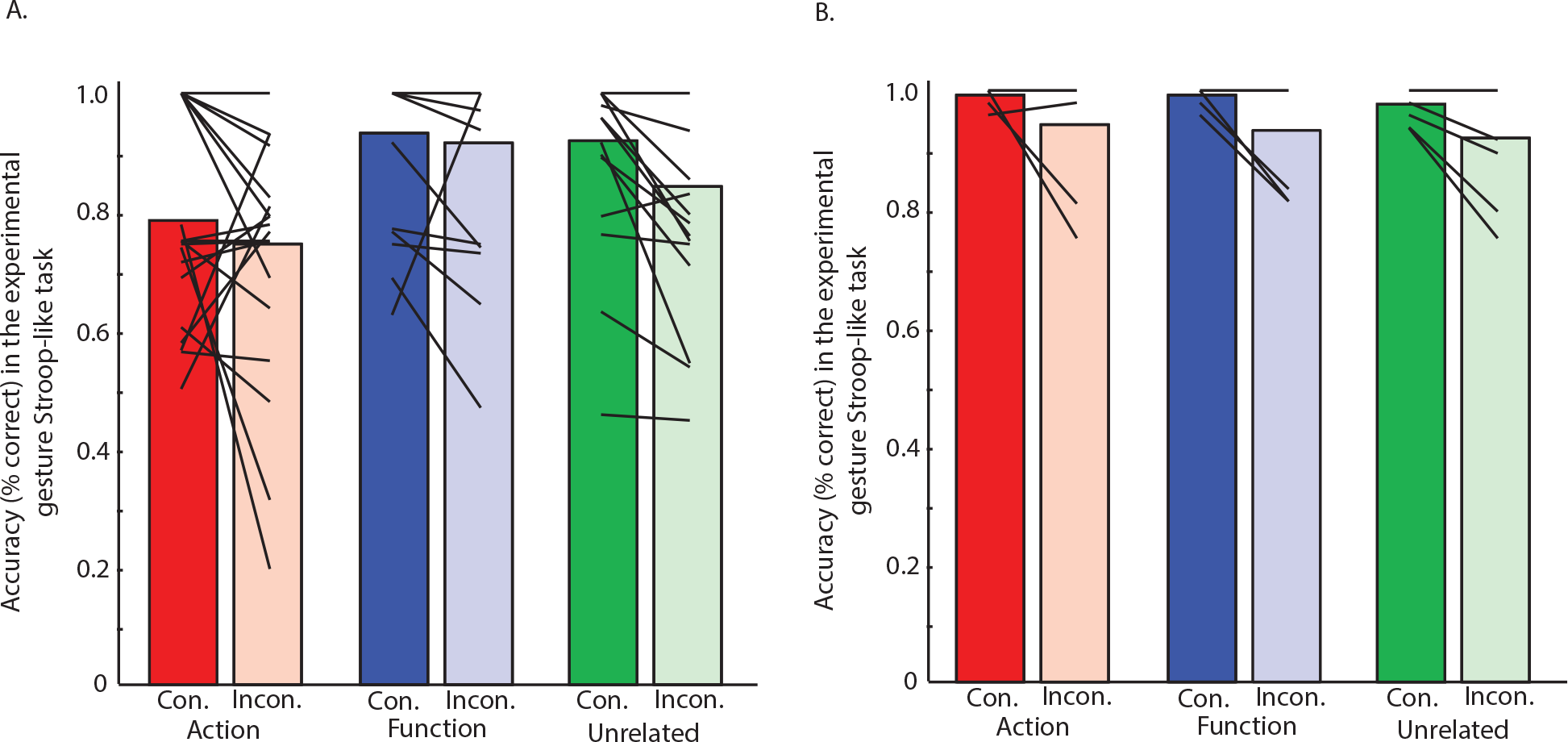
Accuracy in the congruent and incongruent conditions with action, function, and unrelated tool neighbors in the experimental gestural Stroop-like task. Average accuracy for congruent and incongruent trials for **A**) LCVA participants; **B)** neurotypical controls. Note that individual performance is denoted by lines. Abbreviations. Con., congruent; Incon., incongruent.

In Model 1 there was an effect of Congruence (χ^2^ (2) = 83.21, *p* < .001, −.58 ± .07), indicating that participants were more accurate with congruent trials than incongruent trials. There was also an interaction between Group and Congruence (χ^2^ (1) = 23.87, *p* < .001), indicating that neurotypical participants were more accurate than LCVA participants on congruent trials (χ^2^ (1) = 12.28, *p* < .001), 2.38 ± .6) and only marginally more accurate on incongruent trials (χ^2^ (1) = 3.6, *p* = .06), 1.18 ± .6; see Figure 2).

In Model 2 there was an effect of Neighbor type (χ^2^ (2) = 10.47, *p* < .01): Action trials were less accurate than function trials (χ^2^ (1) = 8.09, *p* < .005, −1.36 ± .36) and unrelated trials (χ^2^ (1) = 9.02, *p* < .005, −.76 ± .188), and there was no difference between function and unrelated trials (χ^2^ (1) = 1.79, *p* = .18, −.33 ± .92). Importantly, the Group-by-Neighbor type interaction was also significant (χ^2^ (2) = 35.78, *p* < .001), indicating that control participants were more accurate than LCVA participants on action neighbor trials (χ^2^ (1) = 12.17, *p* < .001, 3.06 ± .08), but were not different on function neighbor trials (χ^2^ (1) = .005, *p* = .93, .17± 2.28) or unrelated neighbor trials (χ^2^ (1) = .6, *p* = .45, .96 ± 1.28). Thus, LCVA participants were deficient in their performance with action neighbors only (see Figure 2). Importantly, there were no significant order effects in either Model (Model 1: χ^2^ (3) = 1.39, *p* = .71; Model 2: χ^2^ (3) = 1.42, *p* = .7).

In Model 3 there was a significant order effect (χ^2^ (5) = 11.92, *p* < .05); however, when entering order as a covariate we found no qualitative change in the effects observed. There was a significant effect of Congruence (χ^2^ (1) = 31.90, *p* < .001, −.046 ± .08), with better performance on congruent trials than incongruent trials, and a significant effect of Neighbor Type, (χ^2^ (2) = 14.87, *p* < .001), indicating that action neighbor trials were less accurate than function (χ^2^ (1) = 11.67, *p* < .001, −1.47 ± .27) and unrelated neighbor trials (χ^2^ (1) = 12.24, *p* < .001, −.9±16). There was no difference between function neighbors and unrelated neighbors (χ^2^ (1) = 2.41, *p* = .14, −.61 ± .35).

Finally, there was an interaction between Neighbor Type and Congruence (χ^2^ (2) = 8.33, *p* < .05), indicating that the congruence effect for action neighbors was significantly reduced in relation to the congruence effect for unrelated (χ^2^ (1) = 5.36, *p* < .05), .44 ± .19) and function neighbors (X^2^ (1) = 8.95, *p* < .01), −.73 ± .24); there was no difference between congruence effects for function and unrelated neighbors (χ^2^ (1) = 0.4, *p* = .53, −.13 ± .21; see Figure 2).

### 3.3 Relationship between Experimental and Grasp-Use Conflict Tasks

Replicating the results from Watson and Buxbaum (2015), data from the test of Grasp-Use Conflict indicated a greater number of hand action errors on conflict (*M*, 0.69; *SD*, 0.11) relative to non-conflict items (*M*, 0.83; *SD*, 0.12), (*t*(20) = 6.10, *p* < .001). In the mixed effect model assessing the relationship between the Experimental and Grasp-Use Conflict tasks, there was an interaction between Grasp-Use Conflict Scores and Neighbor Type (χ^2^ (2) = 19.95, *p* < .001), indicating that the relationship between grasp-use conflict and incongruent action neighbor performance was stronger than the relationship between grasp-use conflict and performance on incongruent function neighbor trials (χ^2^ (1) = 13.99, *p* < .001, 4.64 ± 1.23) and incongruent unrelated trials (χ^2^ (1) = 6.1, *p* < .05, 2.21 ± .89; see Figure 3). There was no difference between the relationship of grasp-use conflict scores and incongruent function neighbor versus incongruent unrelated trials (χ^2^ (1) = .7, *p* = 0.4, 1.21 ± 1.42; see Figure 3). Thus, increased conflict between grasp and use is specifically associated with better performance on incongruent action neighbor trials relative to function and unrelated neighbor types in the experimental task.

**Figure 3.**
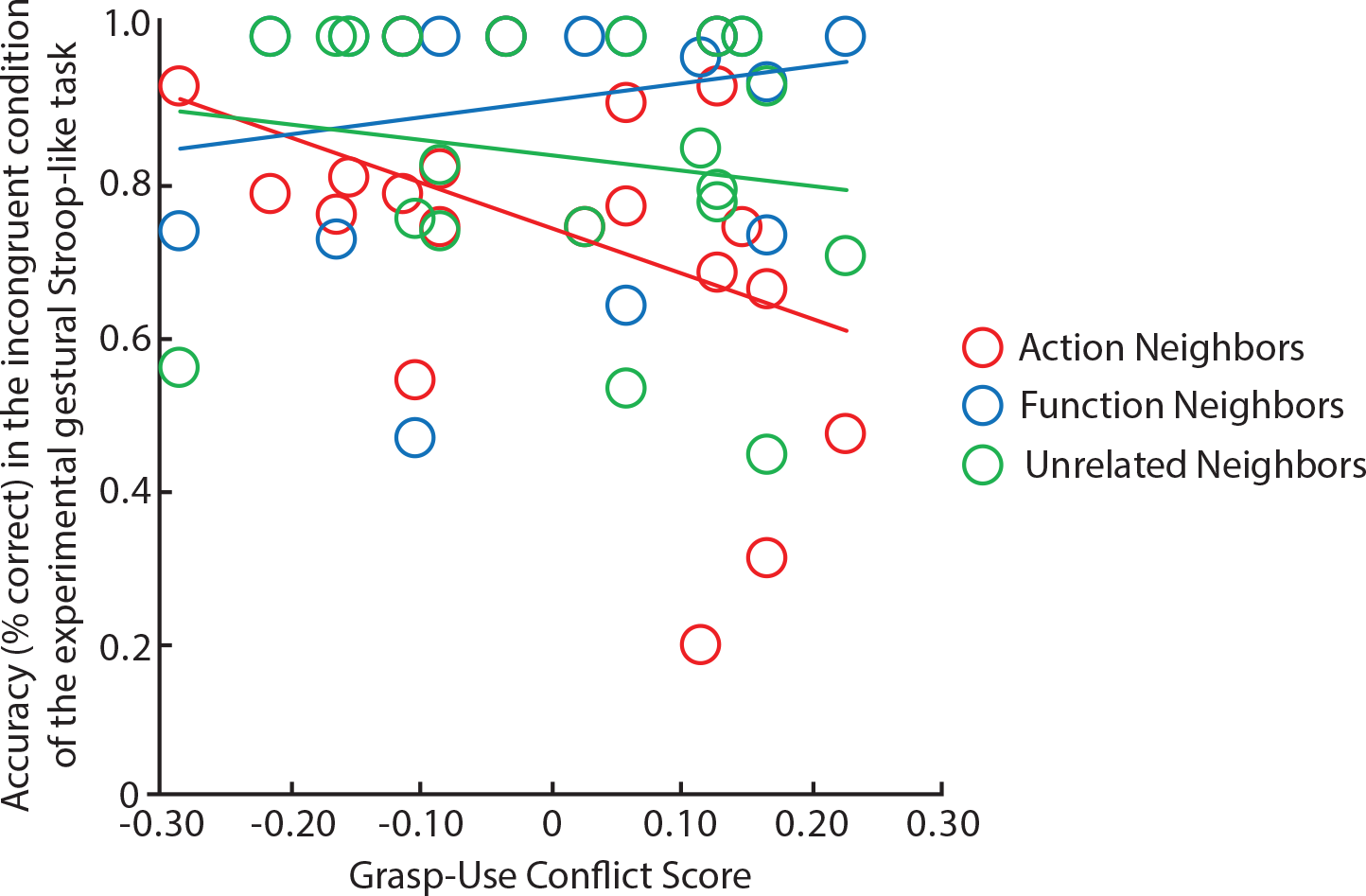
Grasp-use conflict scores plotted against incongruent trials in the experimental gestural Stroop task. Individual LCVA participant data plotted as circles. X-axis: Grasp-Use conflict score. Negative scores indicate higher susceptibility to grasp-use conflict. Y-axis: Incongruent condition of experimental gestural Stroop-like task (% correct).

### 3.4 Support Vector Regression-Lesion Symptom Mapping Results

Figure 4 depicts lesion overlap among the 20 participants with high resolution CT or MRI anatomical data.

**Figure 4.**
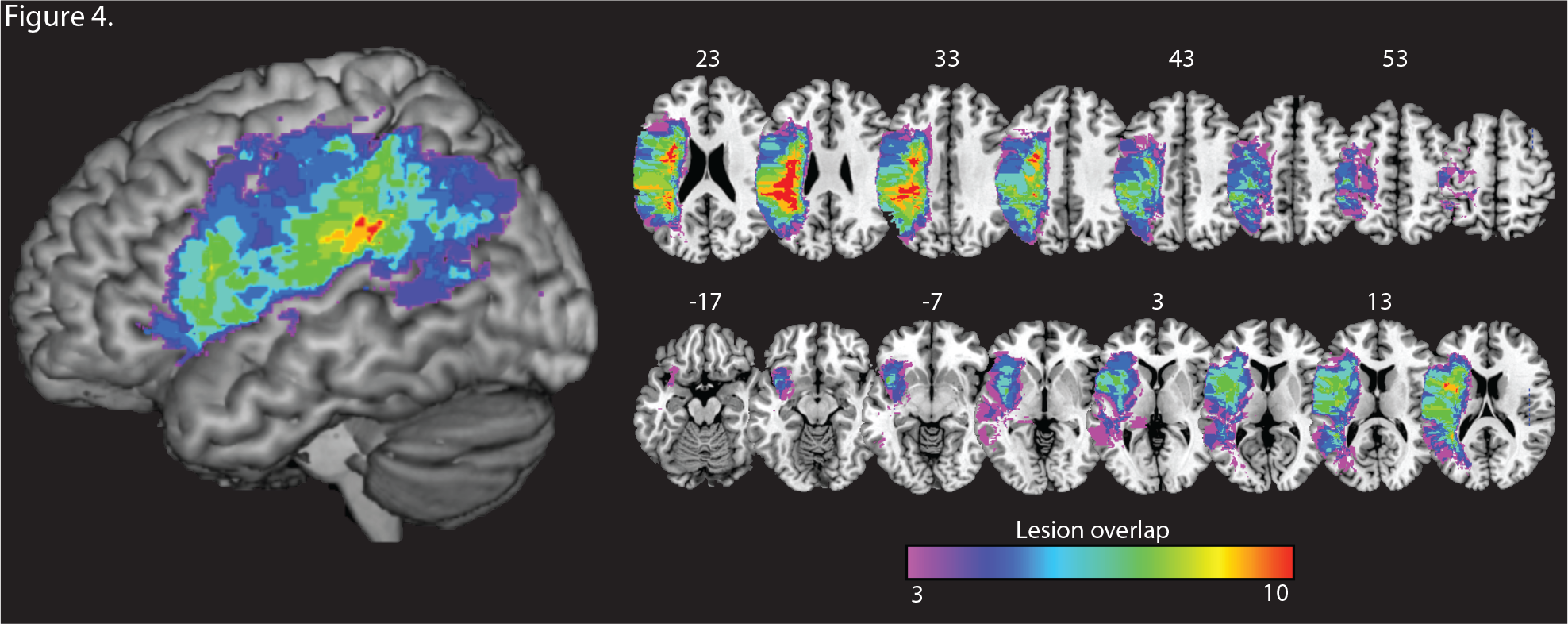
Voxelwise lesion overlap among 20 LCVA participants. Only voxels with at least three participants are included.

Two SVR-LSM analyses were conducted; in the first analysis we identified voxels in which lesions were associated with reduced competition on incongruent action neighbor trials (i.e., hand action performance on incongruent action neighbor trials residualized against hand action performance on congruent action neighbor trials). In the second analysis we identified voxels in which lesions were associated with increased susceptibility to grasp-use competition (i.e., hand action performance when gesturing the use of conflict items residualized against hand action performance with non-conflict items) in the test of grasp-use conflict.

As shown in Figure 5A, lesions to the left SMG, somatosensory and motor cortex (pre- and post-central gyri), and posterior aspect of the left IFG were associated with *better* performance with incongruent action trials in the experimental task; these lesions extended medially to include the left superior longitudinal fasciculus and the left corona radiata as identified by the JHU white matter atlas (see Table 3; see also Supplemental Figure 1). Replicating the voxelwise-lesion symptom mapping results from Watson and Buxbaum (2015), we found that lesions to the left IFG, extending medially into the left anterior insula and posteriorly into the left ventral premotor cortex, as well as subcortically to include aspects of the left superior longitudinal fasciculus, were associated with increased susceptibility to grasp-use conflict (See Figure 5B; see Table 3 for beta values associated with peak MNI coordinates identified in the SVR-LSM analyses).

**Figure 5.**
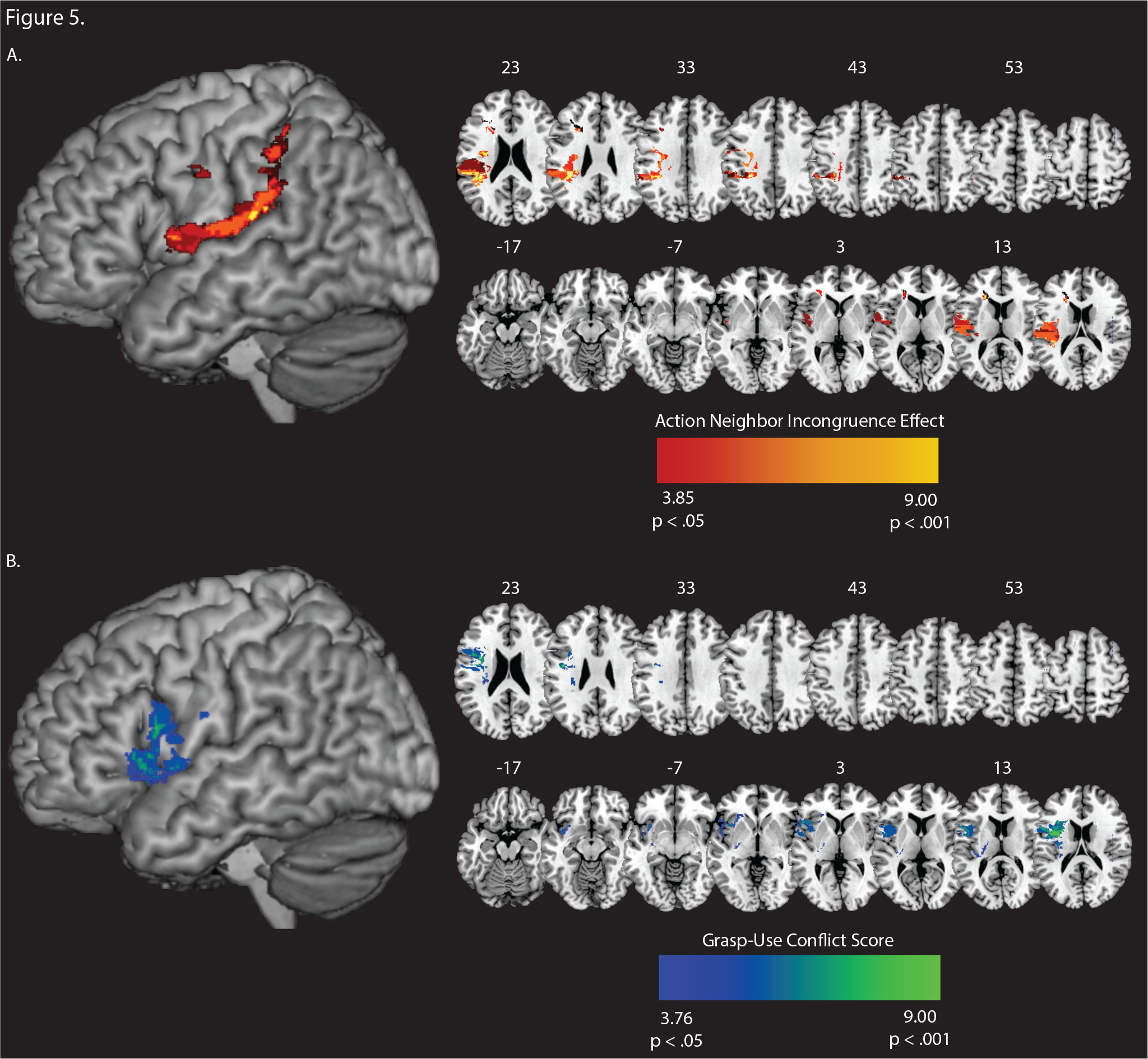
SVR-LSM analyses of data from the experimental gestural Stroop-like task and grasp-use conflict test. **A.** Voxels associated with reduced incongruence effect (incongruent action trials controlling for congruent action trials; red-to-yellow scale) **B.** Voxels associated with grasp-use conflict effect (blue-to-green scale). Whole-brain results are rendered in MNI space in increments of 5 mm. SVR-LSM maps set to a voxelwise threshold of *p* < .05 with 10,000 iterations of a Monte Carlo style permutation analysis; cluster size > 500 contiguous 1mm3 voxels.

**Table 3.**
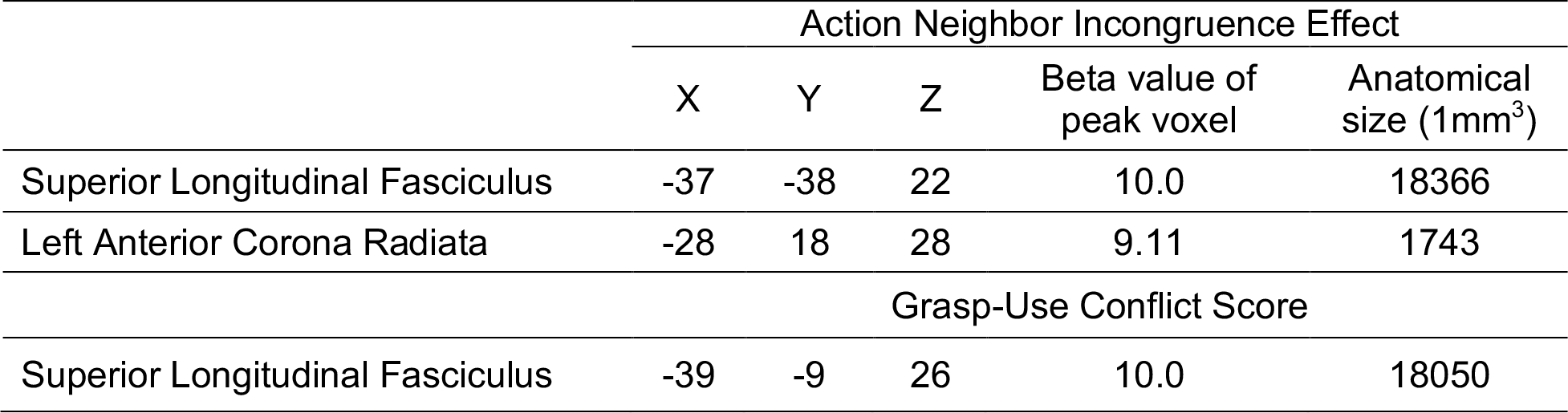
Peak voxels in MNI space identified in the SVR-LSM analysis of the action neighbor incongruence effect from the experimental Stroop-like task, and the grasp-use conflict score. The overlap between peak voxels and major white matter fibers was verified with the Johns Hopkins DTI-based probabilistic white matter tractography atlas.

## 4. General Discussion

This study assessed the hypothesis that reduced automatic activation of use actions in LCVA has specific consequences for patterns of competition within and between tools. We demonstrated that LCVA participants exhibit reduced competition between neighboring tool use actions on a Stroop-like task, as evidenced by a reduced “cost” of producing a gesture associated with a tool’s action neighbor rather than by the depicted tool. Second, we showed that this reduction in competition between tool action neighbors—but not between tool function neighbors—was associated with impaired gesture production in a separate task pitting use against grasp actions within ‘conflict’ tools. Finally, based on SVR-LSM analyses, we also demonstrated that lesions to voxels in the left insula, IFG, SLF, and SMG were associated with reduced competition between tool action neighbors, and that lesions to left IFG and SLF were associated with susceptibility to grasp-use conflict. Together, these data suggest that abnormalities in gesture performance in apraxic individuals are not merely the result of weakened activation of use action representations, but also reflect downstream consequences of weakened activation for competition and action selection. Moreover, they confirm prior observations suggesting that abnormalities in tool action competition are associated with lesions to a specific subset of peri-Sylvian regions in the left hemisphere.

Considerable evidence suggests that the left IFG is critical for performance of a range of semantic control tasks, including naming, word-picture matching, and synonymy judgments with pictures, particularly when items are close neighbors or when demands on control are otherwise high (e.g., Hamilton & Martin, 2005; Mirman & Graziano, 2013; Novick et al., 2009; Nozari, Freund, Breining, Rapp, & Gordon, 2016; Ries et al., 2015; Schnur, Lee, Coslett, Schwartz, & Thompson-Schill, 2005; Schnur et al., 2009; Stampacchia et al., 2018; Thompson-Schill, D’Esposito, Aguirre, & Farah, 1997; Thompson-Schill et al., 1999; Thompson-Schill et al., 1998). Consistent with the proposal that IFG and temporo-parietal regions are nodes of a left hemisphere neural network critical to selection in a broad range of tasks requiring timely and efficient access to semantic information (e.g., Cristescu, Devlin, & Nobre, 2006; Nagel, Schumacher, Goebel, & D’Esposito, 2008; Rodd, Davis, & Johnsrude, 2005)—including action tasks—we have previously suggested that the left SMG briefly buffers candidate actions during the process of selection, whereas the left IFG biases processing such that irrelevant actions are inhibited and task-relevant actions are selected for subsequent production (Buxbaum, 2017; for a related proposal, see Cisek, 2007; Cisek & Kalaska, 2010).

The present results indicate that lesions to partly overlapping brain regions in an IFG-IPL circuit can give rise to abnormalities in both within-tool as well as between-tool action competition. Previous work has demonstrated that grasp information is available more rapidly than use information within single tools; this results in slowing when grasp and use fail to converge on a single response (Jax & Buxbaum, 2010; Lee et al., 2013). From the perspective of classic cognitive winner-take-all models, grasp information rapidly evoked by the shape and size of currently perceived tools must be inhibited in order to select tool use representations (e.g., see Jax & Buxbaum, 2010). Moreover, lesions to the left IFG, insula, and SMG have been associated with abnormal within-tool competition resolution due to increased grasp-on-use conflict (e.g., Jax & Buxbaum, 2013; Watson & Buxbaum, 2015). We did not observe significant SMG voxels associated with within tool (Grasp-Use) conflict in the present study, even at a relaxed threshold (see Supplemental Figure 2). One possibility is that this reflects interesting differences between the portions of the IFG-IPL circuit critical for between- versus within-tool competition resolution (but see Watson & Buxbaum, 2015). Future studies will be required to further explore whether the same regions mediate both aspects of competition.

From a computational perspective, there are at least two non-exclusive interactive activation accounts that may explain how reduced activation of tool use actions can lead to reduced Stroop costs (i.e., improved ability to produce target-incongruent actions). One possibility is that less *precise* activation of tool use actions entails weak activation both of the target action as well as action neighbors, thus facilitating production of the neighbors. Alternatively, weakened automatic activation of use actions may be associated with weaker lateral inhibition of incongruent neighboring actions, rendering them easier to produce. Similar accounts have been proposed in the language domain. For example, a recent computational modeling endeavor suggested that when activation of non-target neighbors is strong, there is a net inhibitory effect on target word selection (in this case, driven in part by lateral inhibition among competing neighbors); in contrast, weak activation of non-target neighbors caused a net facilitative effect on target selection via excitatory connections among the target and non-target neighbors (Chen & Mirman, 2012). On this account, then, weak activation of use representations may lead to net facilitation of the target action relative to its neighbors.

Not all semantic neighbors exert equal effects, as close semantic neighbors will compete to a greater extent than distant semantic neighbors. For example, naming a picture of a dog is frequently slower in the presence of a near semantic neighbor (e.g., cat) than a distant semantic neighbor (e.g., whale) or an unrelated distractor (e.g., couch; see Damian et al., 2001; Fieder et al., 2018; Rose et al., 2018; Vieth et al., 2014; but see Mahon, Costa, Peterson, Vargas, & Caramazza, 2007). The semantic properties that influence neighborhood distance may plausibly include shape, size, color, typical location, or other taxonomic or event-related (thematic) features. In addition, in the tool domain it has become increasingly clear that action is a core semantic feature. Our prior work suggests that tool action similarity takes into account aspects of action appearance that include magnitude of arm movement and posture of the hand (Watson & Buxbaum, 2014), and it is likely that additional features may determine which tools are close in neighborhood space. fMRI experiments in neurotypical adults have reported that tool-related actions may be represented with respect to their visual appearance and motion (Beauchamp, Lee, Haxby, & Martin, 2002), whether they are performed with a tool or the hands (Bracci, Cavina-Pratesi, Ietswaart, Caramazza, & Peelen, 2012; Gallivan, McLean, Valyear, & Culham, 2013), as well as such non-sensory features as the number of sequences they entail (Gallivan, Johnsrude, & Flanagan, 2016) or their abstract function or goal (Wurm & Lingnau, 2015; for review, see Lingnau & Downing, 2015). In future work it will be useful to characterize these and other features in the same tools to explore whether the neighborhood similarity space in the tool action domain is multi-dimensional.

The present data are potentially inconsistent with at least two alternative explanations for action impairments in apraxia following LCVA. First, it has been proposed that technical reasoning, or the ability to derive manipulation information online on the basis of the visual structure of objects, may be the core mechanism disrupted in the apraxia syndrome (e.g., see Osiurak, 2014; Osiurak & Badets, 2016; but see Buxbaum, 2017). Second, it has been argued that apraxics’ apparent deficit in using ‘conflict’ tools is attributable to the greater visual complexity and variability of such objects as compared to non-conflict objects (Bub, Masson, & van Mook, 2018). However, neither of these accounts can accommodate the present data indicating that performance on a Stroop-like gesture task is differentially modulated by neighbor type. Moreover, these accounts cannot explain the observed specific association between reduced interference on the Stroop-like task with action neighbors and impaired performance with conflict tools. Although it is possible that structural affordances, visual complexity, and mechanical problem-solving ability may all influence praxis performance (for discussion, see Buxbaum, 2017; Garcea & Mahon, in press), these alternative explanations alone cannot account for the pattern of abnormal within-item and between-item tool use competition observed here.

There are several limitations of our study worth noting. In our experimental task we used six pairs of tools to study action competition; thus it will be important for future research to identify a larger set of items from which action distance can be derived to study competition among tool action neighbors. Second, although we observed a typical Stroop-like effect for unrelated trials (i.e., better performance with congruent than incongruent items), participants failed to exhibit a typical congruence effect on function trials, a result that was unexpected. Critically however, variability in performance for function neighbors in the experimental task was not related to variability in our measure of grasp-use conflict at the single-item level. Thus, there are competition abnormalities in between-tool actions bearing a specific relationship to competition abnormalities in within-tool actions. On a broader scale, the relatively better performance of LCVA participants on function as compared to action neighborhood trials are consistent with the known dissociation between function and manipulation knowledge in apraxia (Buxbaum & Saffran, 2002; Buxbaum, Veramonti, & Schwartz, 2000; see also Garcea & Mahon, 2012 for dissociations in neurotypical individuals).

### 4.1 Conclusion

In conclusion, the present data complement a number of lines of prior evidence indicating that left hemisphere stroke may be characterized by abnormalities in the competitive processes by which task-appropriate tool action representations are selected from among candidate actions. Although the precise nature of these representations is unclear, they need not be motor plans carrying instructions for the programming of specific muscles (c.f. Oostwoud Wijdenes, Ivry, & Bays, 2016; Wong & Haith, 2017). Rather, we suggest that candidate actions compete in prior processing stages, in which visual action appearance (Buxbaum, 2017), abstract kinematic trajectory (Wong, Haith, & Krakauer, 2015), higher level task goals, and other relatively abstract features determine action similarity (and thus, patterns of competition). Rather than merely reflecting loss of action knowledge, apraxia may be characterized by the consequences of slow and weakened action activation (see Lee et al., 2014) for the downstream selection of appropriate action representations that inform subsequent motor planning.

## Acknowledgements

We would like to thank Cortney Howard, Leyla Tarhan, and Louisa Smith for coding participants’ gestures; Christine Watson for sharing her skills and tool norming data; and H. Branch Coslett for help with lesion segmentation. This research was supported by NIH grant R01 NS099061 to L.J. Buxbaum. Preparation of this manuscript was supported by a Moss Rehabilitation Research Institute/University of Pennsylvania postdoctoral training fellowship (NIH 5T32HD071844-05).

## Author Information

The authors declare no competing financial interests. Correspondence should be addressed to F.E. Garcea.

## Supplemental Online Materials

**Supplementary Table 1.**
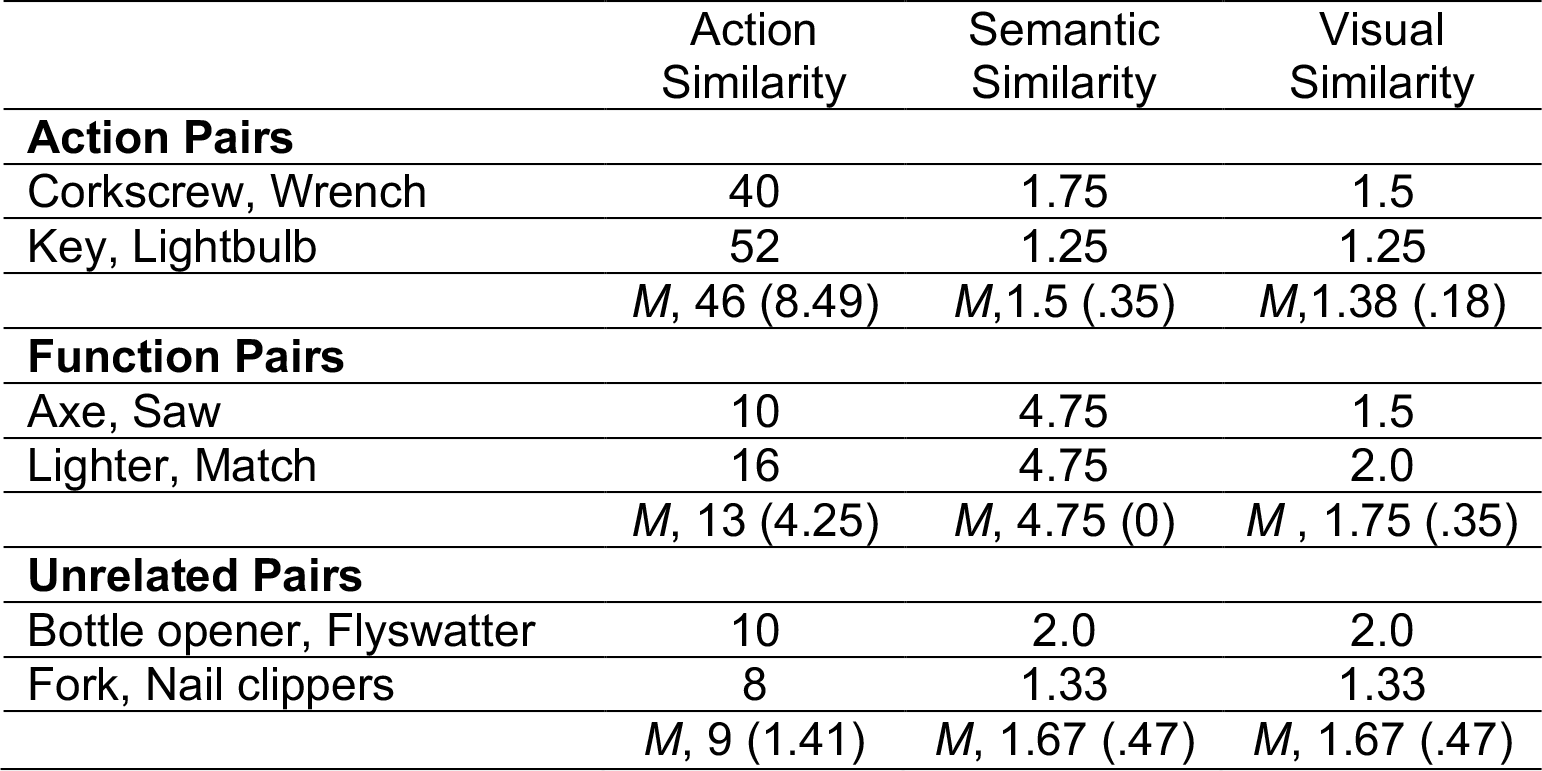
Mean (and standard deviation) ratings of action, semantic, and visual similarity (from Watson and Buxbaum, 2014) for tool pairs used in the experimental gestural Stroop-like task. Maximum action similarity = 72; maximum semantic and visual similarity = 5; see Watson and Buxbaum (2014) for details.

**Supplementary Table 2.**
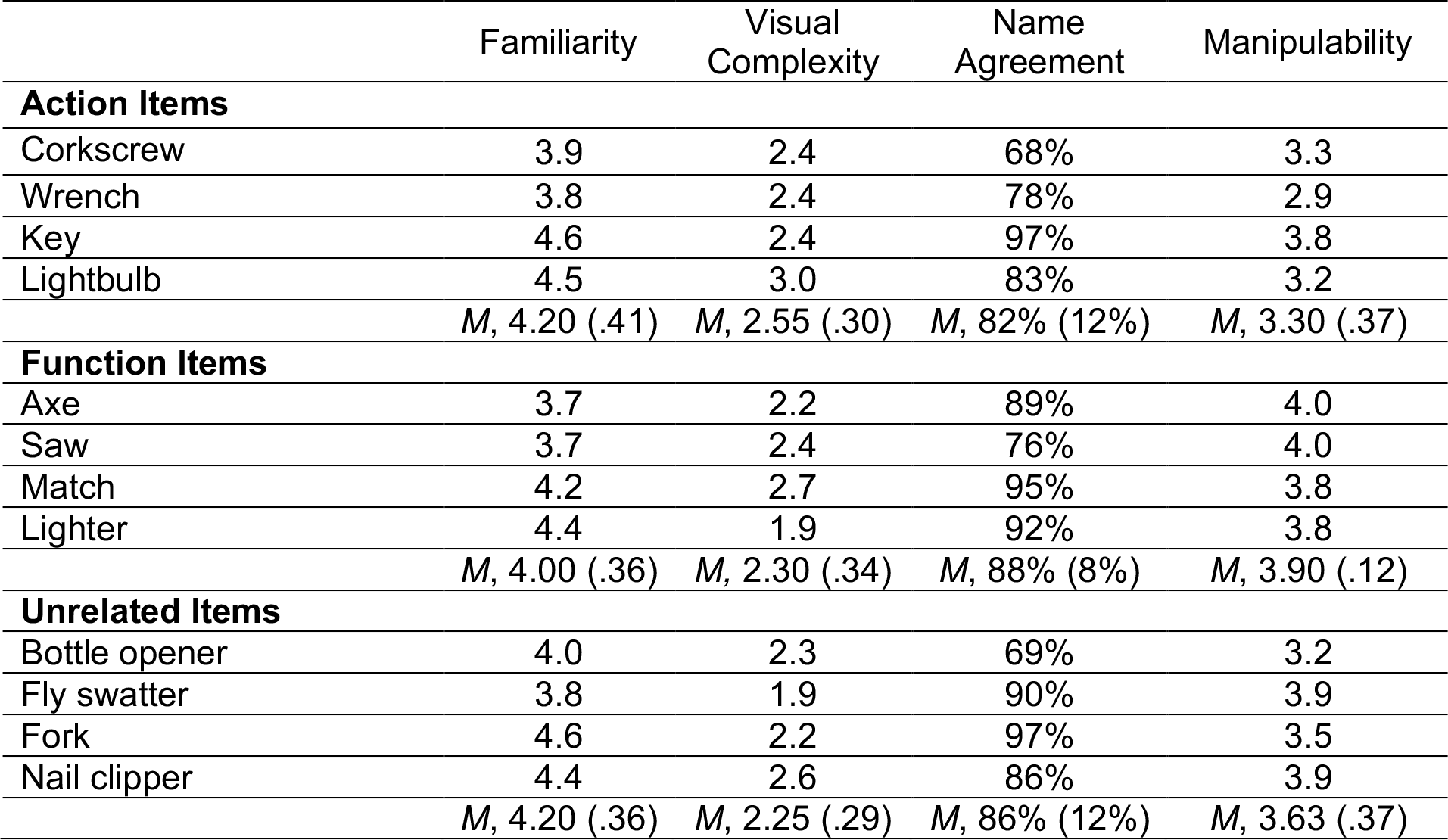
Mean (and standard deviation) ratings of familiarity, visual complexity, name agreement, and degree of manipulability for tools used in the experimental gestural Stroop-like task, derived from the BOSS dataset (Brodeur et al., 2010). See Brodeur et al (2010) for details.

**Supplementary Figure 1.**
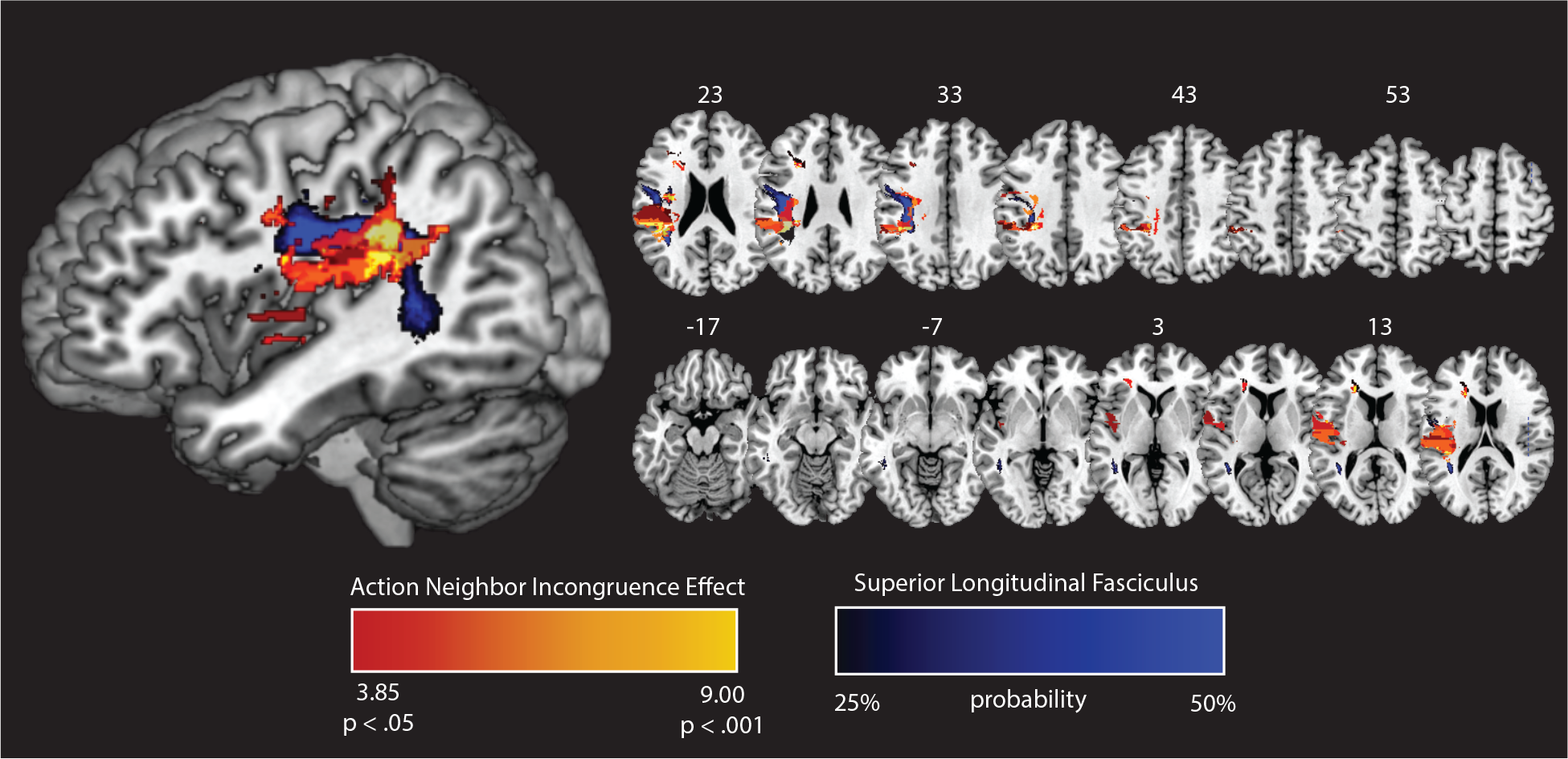
Overlap between the action neighbor incongruence effect in the experimental gestural Stroop-like task and the left superior longitudinal fasciculus defined by the Johns Hopkins White Matter Atlas in FSL.

**Supplementary Figure 2.**
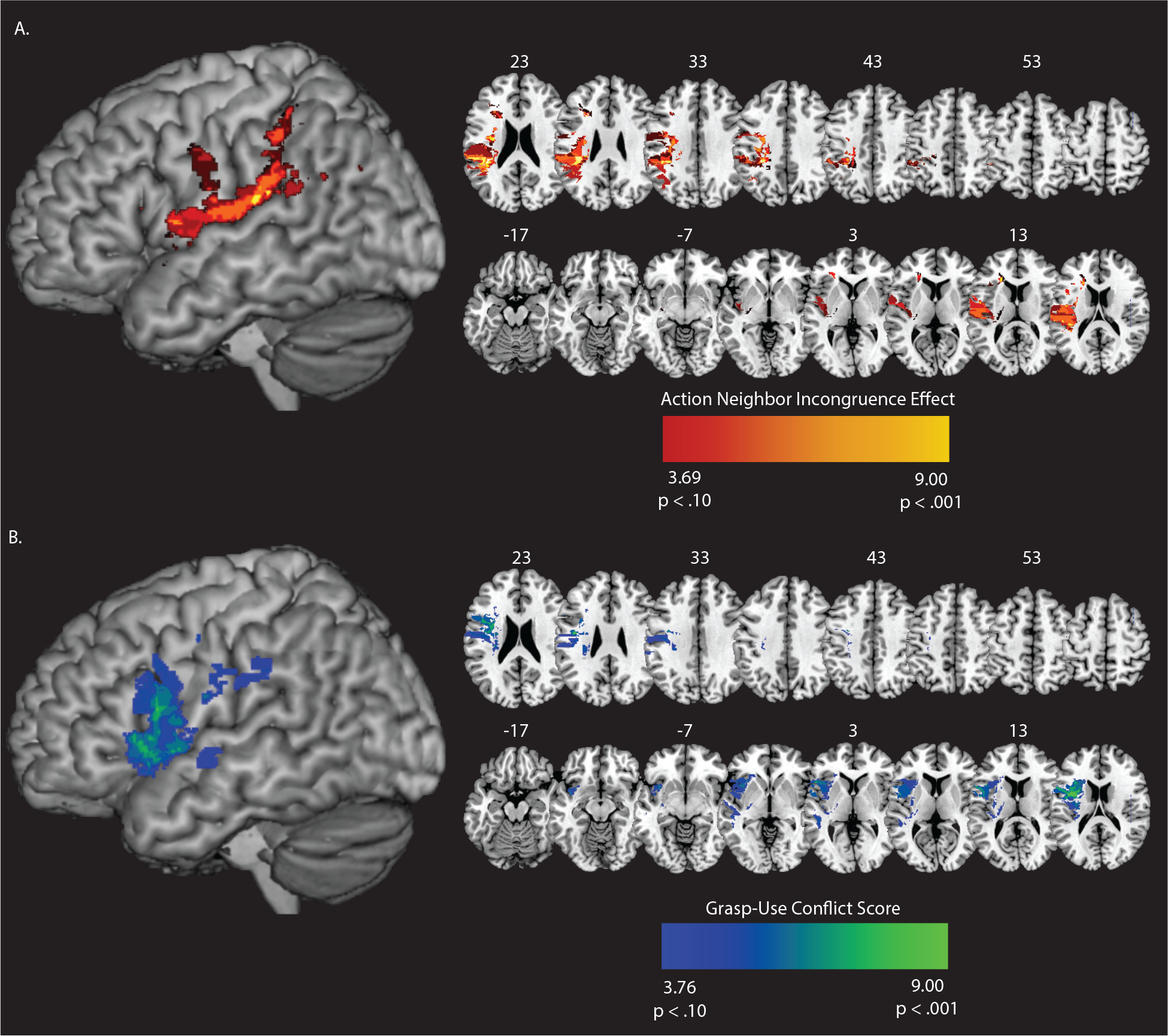
Support vector regression lesion-symptom mapping results at a relaxed alpha threshold (*p* < .10). **A.** The action neighbor incongruence effect from the experimental gestural Stroop-like task. **B.** The grasp-use conflict score from the test of grasp-use conflict. Voxels identified in both analyses survive a Monte Carlo style voxelwise permutation analysis (10,000 iterations), and form a cluster of at least 500 contiguous voxels.

## Footnotes

1 We use the term “hand action” and “hand posture” interchangeably throughout the text.

